# Hybrid RY-mers increase aDNA alignment sensitivity

**DOI:** 10.64898/2026.11.05.764514

**Authors:** Ivan Tolstoganov, Kristoffer Sahlin

## Abstract

One of the key steps in ancient DNA data analysis is the accurate identification of endogenous short reads that originate from the ancient genome. Currently, this step is performed using general-purpose sequence aligners, such as BWA-aln, with parameter settings optimized for sensitivity. This approach results in increased runtime, even relative to standard DNA alignment workflows, and is not sufficient for aligning samples that are highly divergent from a reference database. We introduce hybrid RY-mers, a seed construct designed to incorporate aDNA-specific sequence damage. We adapt hybrid RY-mers to the existing concept of multi-context seeds and implement them in the read aligner strobealign. On simulated genomic and metagenomic datasets, we show that hybrid RY-mers result in higher mapping accuracy, while also reducing runtime by 4-6 times compared to the state-of-the-art aDNA alignment method BWA-aln. Furthermore, substituting aDNA aligner MALT in the metagenomic tool-suite aMeta with our new alignment approach demonstrated that aMeta gave a faster and less memory consuming detection of ancient taxa present in the samples.

## Introduction

Ancient DNA (aDNA) sequencing provides evidence of past organisms and microbial communities [1, 2]. As with most analyses of sequencing data, sequence alignment is a key step in aDNA analysis, required for variant calling, taxonomic profiling, and authentication of ancient microbial DNA.

However, sequence alignment of aDNA samples is particularly complicated by several factors. First, cytosine deamination produces uracil, read as thymine, leading to apparent C-to-T and complementary G-to-A substitutions concentrated near fragment ends in double-stranded libraries [3]. Second, DNA degradation results in short reads with few seeds for locating candidate alignments. Finally, samples can contain DNA from the original organism and its microbes, organisms that colonized remains after death, and contamination introduced during excavation or laboratory handling [4].

General-purpose aligners such as BWA-aln [5], BWA-MEM [6] and Bowtie2 [7] can accommodate damaged reads through parameter adjustments, although alignnment sensitivity depends on read length and search settings [8]. The metagenomic aligner MALT [9], which is widely used in aDNA studies as part of the pathogen-screening pipeline HOPS [10], combines spaced-seed alignment [11] with taxonomic assignment. Spaced seeds can match over substitutions by excluding selected positions from comparison and replacing them with a wildcard character. However, the wildcard characters are not specific to deamination, since any substitution is accepted at an ignored position. RY-mer constructs introduced in the SAFARI aligner [12] use a more specific rule by encoding each nucleotide as a purine (R; A/G) or pyrimidine (Y; C/T) [13]. Because C-to-T and G-to-A substitutions preserve these classes, RY-mers retain matches across deaminated bases. R/Y-encoding is also used for mapping bisulfide sequencing reads [14]. Earlier transition-constrained seeds [15] can be viewed as an extension of RY-mers, allowing R/Y encoding or a spaced-seed wildcard at arbitrary positions of the seed.

While being more aDNA-specific than spaced seeds, RY-mers still discard nucleotide distinctions at undamaged positions. Assuming independent, equally frequent nucleotides, an RY-mer of length *k* carries *k* bits of information, compared with 2*k* bits for a nucleotide seed of the same length. A lower amount of seed information increases the number of spurious matches requiring filtering and alignment, particularly in large or repetitive references. These shortcomings motivate retaining nucleotide identity where possible and relaxing matches where damage is expected.

To this end, we develop *hybrid RY-mers*, a combination of RY-mer and *multi-context seed* [16] concepts. Multi-context seeds (MCS) encode strobemers [17] and their constituent *k*-mers in a single hash value, allowing both to be searched in the same index. Strobemers are longer, more specific, or *full* seeds, while *k*-mers are shorter and less specific *partial* seeds. A hybrid RY-mer follows the same principle as MCS by storing the full seed in the hash along with its less specific part. Here, the full seed is a *k*-mer, while the partial seed is a string of *k* characters, each character encoded either in the *{A, C, G, T }* or *{R, Y }* alphabet. This structure of the partial seed allows us to rescue the reads that do not have any *k*-mer matches due to deamination damage.

We implemented strobealign-HRY, a version of strobealign [18] that uses hybrid RY-mers instead of multi-context seeds. This version, denoted as strobealign-HRY, is used as the ancient DNA alignment mode for strobealign. We benchmark strobealign-HRY against BWA-mem [6], BWA-aln [5], Bowtie2 [7], and MALT [9] on datasets with simulated deaminated damage, showing that strobealign-HRY results in higher mapping accuracy than commonly used aDNA aligners.

## Methods

### Multi-context seeds

A strobemer is a seed formed by linking shorter k-mers, called strobes, sampled at variable distances along a sequence [17]. Originally, multi-context seeds (MCS) were introduced to search for a strobemer or a prefix of its constituent strobes in the same index [16]. Here, we extend the definition to bit strings representing arbitrary seeds or their hash values.

Let *s* be a bit string and let (*I*_1_, …, *I*_*n*_) be an ordered partition of its positions into nonempty, disjoint subsequences. For each *i*, let *s*_*i*_ = *s*[*I*_*i*_] be the subsequence containing the bits at positions in *I*_*i*_. Let *h*_*L*_ map bit strings to *L*-bit hashes, and let 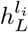 denote the *l*_*i*_-bit prefix of such a hash. Given positive integers *l*_1_, …, *l*_*n*_ with 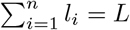, we define the corresponding *n-context seed*, or *n*-MCS, as

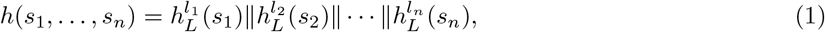

where *∥* denotes concatenation. The first *l*_1_ bits form the *base hash*, and the remaining components are auxiliary hashes. For instance, given *n* bit-strings representing 2-bit nucleotide encodings of *n* DNA strings *s*_*i*_ (e.g., strobes), each bit-string is hashed into *L*-bit hashes and concatenated.

Two seeds have a *full match* if their complete MCS hashes are identical. They have a *partial match* at level *j < n* if the prefixes containing their first *j* component hashes are identical (see Figure 1b). The component order determines which seed subsets can be queried.

**Fig 1.**
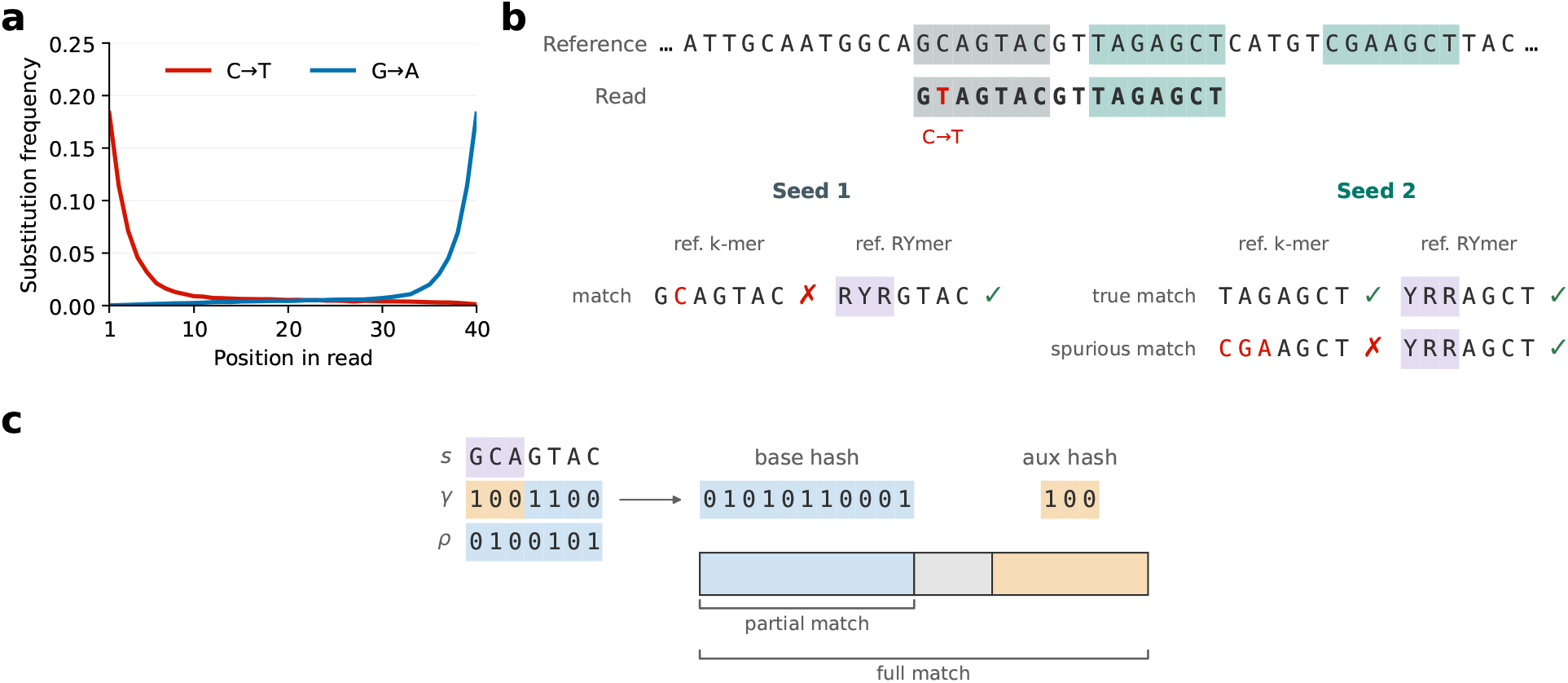
Overview of hybrid RY-mers. a. C-to-T and G-to-A substitution frequencies along simulated 40 bp reads, using the default gargammel simulation parameters. b. Seeding with hybrid RY-mers (*k* = 7, *ℓ* = 3). Violet shaded positions are R/Y-encoded. Seed 1 covers a deaminated base with its R/Y prefix. The corresponding *k*-mer has no exact match in the reference, but partial matched is obtained using RY-mer prefix. Seed 2 has a full match at its true location and a partial match at a spurious location, which is not used since the full lookup succeeds. c. Hash layout of a hybrid RY-mer for *s* = GCAGTAC. The R/Y-encoded prefix and the nucleotide suffix produce the base hash, and the remaining bits of the prefix produce the auxiliary hash by applying hash function. The orientation bit (grey) is stored between the two components. Partial matches compare the base hash, and full matches compare the entire hash.

### Hybrid RY-mers

We define a hybrid RY-mer as a 2-MCS consisting of two parts, with the input string *s* corresponding to a bit representation of a *k*-mer. The base hash *B*_*P*_ represents a *k*-mer with some characters replaced with R/Y according to the pattern *P* . The rest of the hash *A*_*P*_ consists of the bits required to restore the original *k*-mer representation.

Formally, let *s ∈ {A, C, G, T }*^*k*^ be a *k*-mer and let a nonempty set *P ⊆ {*0, …, *k −* 1*}* specify its R/Y positions. We define two binary encodings of nucleotides,

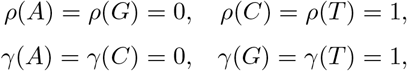

and extend both encodings to strings by concatenation. The pair *γ*(*a*)*ρ*(*a*) uniquely encodes each nucleotide *a*. For example, the encoding of nucleotide A is *γ*(*A*)*ρ*(*A*) = 00 and G is *γ*(*G*)*ρ*(*G*) = 01. Note that *ρ* is here the R/Y encoding [13], (maps A and G to 0 and C and T to 1).

For the pattern *P*, define the mixed-alphabet component *B*_*P*_ (*s*) and its complementary component *A*_*P*_ (*s*) as

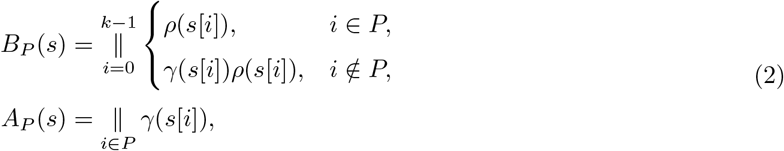

where positions are traversed in increasing order. These components partition the two-bit encoding of *s* and have lengths 2*k −* |*P* | and |*P* |. Figure 1c visualizes the layout.

Given a hash function *h*_*L*_ and positive hash widths *l*_1_, *l*_2_ with *l*_1_ + *l*_2_ = *L*, the *hybrid RY-mer* is the 2-MCS

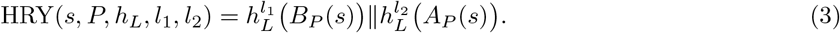

The first component hash is used for partial matches. In the absence of hash collisions, a full match requires nucleotide identity throughout *s*, while a partial match requires R/Y identity at positions in *P* and nucleotide identity elsewhere. Partial matches tolerate transitions at R/Y positions, as well as the C-to-T and G-to-A substitutions resulting from deamination damage.

The above formulation allows specifying R/Y and nucleotide identity requirements at general positions in the seed, enabling various patterns (similar to spaced seeds). However, since cytosine deamination is concentrated near fragment ends [3], we use the *prefix layout P*_*ℓ*_ = *{*0, …, *ℓ −* 1*}*, with 1 *≤ ℓ ≤ k*, to require only R/Y identity at the *ℓ* prefix positions of a seed.

### Implementation

We implement hybrid RY-mers as part of strobealign, as an ancient DNA mode (strobealign-HRY), but otherwise generally following the MCS indexing and lookup approach [16].

*Seed selection* In order to preserve matches in R/Y encoding between forward- and reverse-oriented strings, we decide the canonicity of a *k*-mer based on its *ρ*-encoding, rather than full *k*-mer hash. Specifically, for a nucleotide string *u*, let *u* denote its reverse complement and define *K*(*u*) = (*ρ*(*u*), *γ*(*u*)). We compare these pairs lexicographically, first by the R/Y bit string and then by the complementary bit string. We define *canonical representation* as

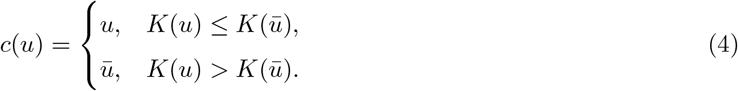

We encode the *canonical orientation* as *o*(*u*) = 1 when the forward representation is selected and *o*(*u*) = 0 otherwise. Equal pairs select the forward representation. Orientation is determined before hashing and independently of the pattern *P* .

Open syncmers subsample *k*-mers according to the position of a minimum-hash substring [19]. For a substring length *q* and a zero-based offset *t*, where 1 *≤ q ≤ k* and 0 *≤ t ≤ k − q*, a *k*-mer *s* is selected iff

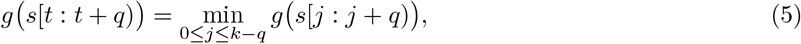

where *s*[*j* : *j* + *q*) denotes the substring of length *q* starting at position *j*. Our implementation uses *g*(*u*) = *h*_sel_(*ρ*(*c*(*u*))), with a separate hash function *h*_sel_ for selection. We require *k − q* to be even and set *t* = (*k − q*)*/*2, which makes selection invariant under reverse complementation. Each selected *k*-mer produces HRY(*c*(*s*), *P*_*ℓ*_, *h*_*L*_, *l*_1_, *l*_2_) and its orientation bit *o*(*s*).

Note that the syncmers are selected on the R/Y-encoded s-mers, i.e., *g*(*u*) = *h*_sel_(*ρ*(*c*(*u*))), rather than *g*(*u*) = *h*_sel_(*c*(*u*)). As a result, a deamination event in a read does not change which positions are selected as syncmers, since C*→*T and G*→*A substitutions leave the R/Y encoding unchanged. Had we selected syncmers in nucleotide space, a deamination event would likely alter the hash values of the s-mers overlapping it and thereby the set of selected syncmers, so the same syncmers would rarely be selected in the reference and the read, even if they were subsequently R/Y-encoded.

*Indexing layout* Reference seeds are stored in an array sorted by hash value, with a prefix lookup table locating the range to search [20]. Each entry contains a hybrid RY-mer hash, a reference identifier, and a reference position. The base and auxiliary hashes represent 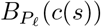 and 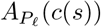, respectively. Matching the base hash identifies a contiguous range of candidate partial matches in the index.

The main layout change compared to strobealign v0.16 version described in [16] is that we store *o*(*s*) in a bit between the base and auxiliary hashes. Orientation-aware searches include this bit, while a base-hash lookup can mask it out to retrieve both orientations. This orientation bit is used in both HRY and default modes of strobealign.

*Querying* Under the default MCS search strategy, we first query the full hash. If it is absent in the index, we query the base hash for a partial match. In ancient DNA mode, partial lookup is restricted to k-mers that start within the first *ℓ* positions of the read or end within its last *ℓ* positions. For reverse-complement queries, we reuse the hybrid RY-mer hash, reverse the seed coordinates, and flip the stored orientation bit. Repetitive hits (*>* 1000 occurrences by default) are filtered before chaining. Seed matches are chained and extended using the original nucleotide sequences and strobealign’s alignment procedure [18].

## Results

### Benchmark setup

We evaluate Strobealign-HRY on simulated ancient DNA reads with known ground truth, measuring mapping accuracy, runtime, and peak memory usage. We assess endogenous-read recovery across mutation rates and separately evaluate reads carrying simulated deamination damage. We compare strobealign-HRY with BWA-aln (v.0.7.19), BWA-MEM (v.0.7.19), Bowtie2 (v. 2.5.0), and a standard version of strobealign, but with parameters tuned for aDNA alignment. We denote it strobealign-DK, for dynamic k-mers, as it uses multi-context seeds [16] without a gap between strobes [17]. Tool settings and timing definitions are provided in Supplementary Section S1.

We use Mason [21] variator to generate substitutions at rates of 0.5%, 2% and 5%, with corresponding small-indel rates of 0.1%, 0.2% and 0.5%. The maximum indel size is 100 bp. Reads are simulated from the original sequences and mapped against the modified references. Ground-truth coordinates are transferred to the modified references using g2gtools https://github.com/churchill-lab/g2gtools.

An endogenous read is considered to be correctly mapped when its primary alignment passes the alignment-score threshold and overlaps its ground-truth interval on the correct reference sequence. Accuracy is the percentage of all retained endogenous reads mapped correctly, including unmapped reads in the denominator. Mapping rate includes both correct and incorrect placements that pass the threshold. Reads without an accepted alignment are counted as unmapped or filtered. We calculate these measures separately for reads tagged by the simulator as carrying deamination. For the comparison between deaminated and non-deaminated reads, accuracy is calculated within each subset. We only retain reads with an alignment score exceeding 60. Alignment scores for all aligners are computed using strobealign’s scoring scheme [18]. Secondary and supplementary alignments are excluded.

### Simulated datasets

The benchmark uses the fruit fly genome, CHM13 assembly [22], and reference genomes from the ZymoBIOMICS Microbial Community Standard [23], each modified at different mutation rates. For each dataset, gargammel [24] simulates 500,000 fragments of 40 bp, comprising 80% endogenous material and 20% bacterial background from the k14 microbial community [25]. See Figure 1a for the deamination profile. Here, endogenous denotes sequences sampled from the target reference set. Both endogenous and background fragments undergo simulated damage, using the default gargammel deamination damage simulation parameters [24]. Reads undergo adapter trimming and quality filtering with fastp.

### Read mapping evaluation

#### Mapping accuracy

Strobealign-HRY has the highest mapping accuracy in eight of the nine datasets (Figure 2a). The exception is CHM13 at 2% substitutions, where its accuracy is *∼*0.1 percentage points below that of BWA-aln. However, at this substitution level, BWA-aln maps endogenous reads at a significantly lower precision. BWA-aln maps 91.2% of endogenous reads, compared with 86.3% for Strobealign-HRY, where incorrect placements account for 20.42% and 15.67% of endogenous reads, respectively (Figure 2a). In every dataset, Strobealign-HRY accuracy exceeds that of Strobealign-DK.

**Fig 2.**
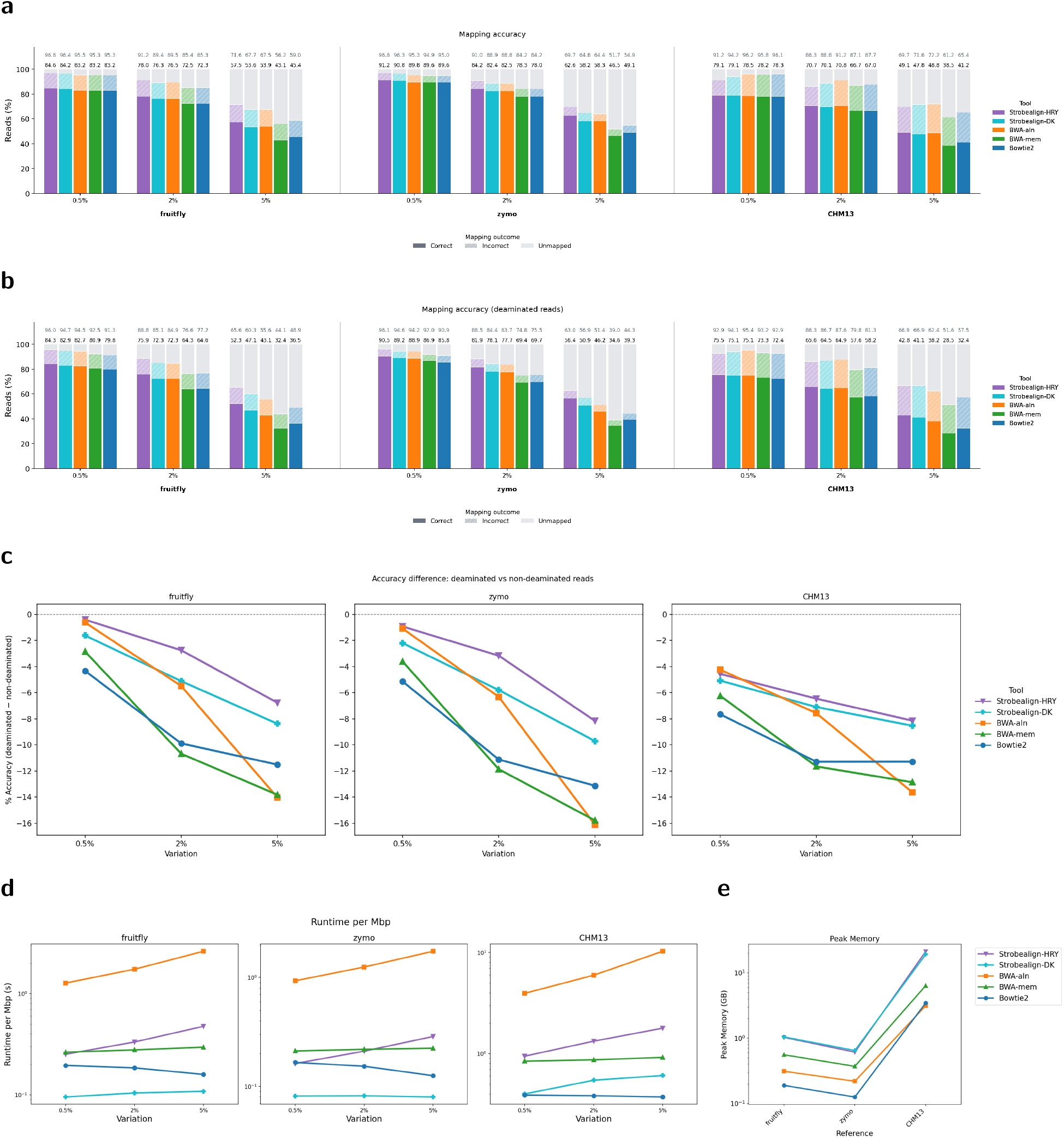
Read mapping evaluation across fruit fly, Zymo and CHM13 references at substitution rates of 0.5%, 2% and 5%. a. Mapping outcomes for all endogenous reads. Solid segments show correctly mapped reads, hatched segments show incorrectly mapped reads, and grey segments show reads without an alignment passing the alignment score threshold. The lower and upper numerical labels above each bar report accuracy and mapping rate, respectively. b. Read mapping outcomes evaluated for reads carrying simulated deamination. c. Accuracy on deaminated reads minus accuracy on non-deaminated reads, in percentage points. d. Mapping runtime in seconds per megabase of simulated read sequence (log scale) e. Peak resident memory in GiB (log scale). For every reference, mean values over variation rates are shown.

#### Deaminated-specific accuracy

Reads carrying simulated deamination constitute 21.84-25.07% of endogenous reads across datasets (mean 23.20%). Strobealign-HRY has the highest accuracy on this subset in all nine datasets (Figure 2b), showing that hybrid RY-mers perform better over deaminated regions. At 5% substitutions, its deaminated-read accuracy is 52.27% for fruit fly, 56.44% for Zymo, and 42.79% for CHM13. Every aligner has lower accuracy on deaminated reads than on non-deaminated reads (Figure 2c), which we denote as *accuracy deficit*. Strobealign-HRY has the smallest accuracy deficit in eight datasets, while BWA-aln has a slightly smaller accuracy deficit for CHM13 at 0.5% substitutions.

#### Runtime and memory usage

BWA-aln has the longest mapping runtime in every dataset, taking 4-6 times as long as Strobealign-HRY (Figure 2d). Strobealign-HRY takes 2-4.3 times as long as the non-aDNA specific strobealign-DK. Strobealign-DK is fastest for fruit fly and Zymo, whereas Bowtie2 is fastest for CHM13. Both strobealign configurations use more memory than the other aligners across all datasets (Figure 2e). Bowtie2 uses the least memory for fruit fly and Zymo, while BWA-aln uses the least for CHM13.

#### Seed size

Strobealign-HRY’s higher accuracy, longer runtime, and greater memory usage than strobealign-DK are partly explained by its smaller effective seed size compared with Strobealign-DK and BWA-MEM. Strobealign-HRY uses hybrid RY-mers with k = 17 and l = 8 by default, equivalent to 13-mers in seed entropy. To investigate the effect of seed size on accuracy and runtime, we compare BWA-MEM, strobealign-HR and strobealign-DK across k values from 13 to 21. Reducing k below 17 increases accuracy by less than 2% (Supplementary Figure S1), while substantially increasing runtime (Supplementary Figure S2).

### Metagenomic benchmark

We next evaluated how the choice of aligner affects microbial detection and authentication in ancient metagenomic samples. We used aMeta, an established workflow that combines taxonomic pre-screening, sequence alignment, and validation of microbial presence and ancient origin [26]. Its separation of pre-screening from alignment allows us to compare alignment methods using the same microbial candidates and reference sequences.

The benchmark follows the simulation design of [26], with ten samples drawn from 18 ancient and 17 modern microbial species, together with human DNA. Each sample contains 500,000 single-end reads, obtained by combining the first mates from two gargammel simulations of 250,000 fragments each. The microbial fraction varies from 30% to 70% across samples. Ancient microbial fragments follow a log-normal length distribution with a median of approximately 42 bp and undergo simulated deamination using the Briggs model [3]; modern microbial fragments follow an empirical length distribution without deamination. Both simulations include sequencing errors and generate 125 bp reads before adapter trimming. Taxon identities and ancient or modern status are recovered from the simulated reads to establish the ground truth.

Following aMeta workflow, we run KrakenUniq to pre-screen the data [27, 28], using the prebuilt Standard database and aMeta’s filtering thresholds of 1,000 unique k-mers and 200 assigned reads per taxon. We then compare aMeta’s MALT alignment and taxonomic binning step with Strobealign-HRY, BWA-aln, BWA-mem, or Bowtie2 followed by ngsLCA [29]. MALT performs read alignment, followed by taxonomic assignment of reads using the lowest common ancestor (LCA) of reference taxa with at least one secondary alignment [9]. For the other benchmarked aligners, the aligner produces the candidate placements, and ngsLCA performs the taxonomic assignment. The resulting assignments are evaluated using aMeta authentication scoring. To every candidate species, aMeta assigns 9 scores ranging from 1 to 2 (total 10). The individual scores evaluate read length distribution, coverage depth, coverage breadth, likelihood of ancient origin based on the deamination damage profile, ANI with the database reference genome, and edit distance distribution.

We evaluate the resulting aMeta authentication scores against the simulated ancient microbial community. For each authentication score threshold *t*, taxa scoring at least *t* are predicted to be ancient, and their agreement with the true ancient set is measured by Jaccard similarity, averaged across the ten samples. To isolate the post-screening comparison, both predictions and ground truth are restricted to taxa retained by KrakenUniq. These KrakenUniq-adjusted results therefore do not count microbes missed during pre-screening as alignment or authentication failures. Human sequences are excluded from the evaluation.

The choice of the alignment method has little effect on the recovery of the ancient community at score thresholds of 7 or 8 (Figure 3a). At *t* = 7, all five methods recover 66 of the 72 ancient taxon-sample combinations, with no false positives and an identical mean Jaccard similarity of 0.91. At *t* = 6, however, the methods produce five to nine false positives, since some of the aMeta authentication criteria, such as evenness of coverage and edit distance monotonicity, are also applicable to modern taxa. Therefore, threshold values of 6 or lower are less applicable to ancient species set recovery. Thresholds of 8, 9, or 10 reject more true ancient taxa and reduce agreement with the ground truth. Results for individual authentication criteria are reported in Supplementary Section S2 and Supplementary Figure S4.

**Fig 3.**
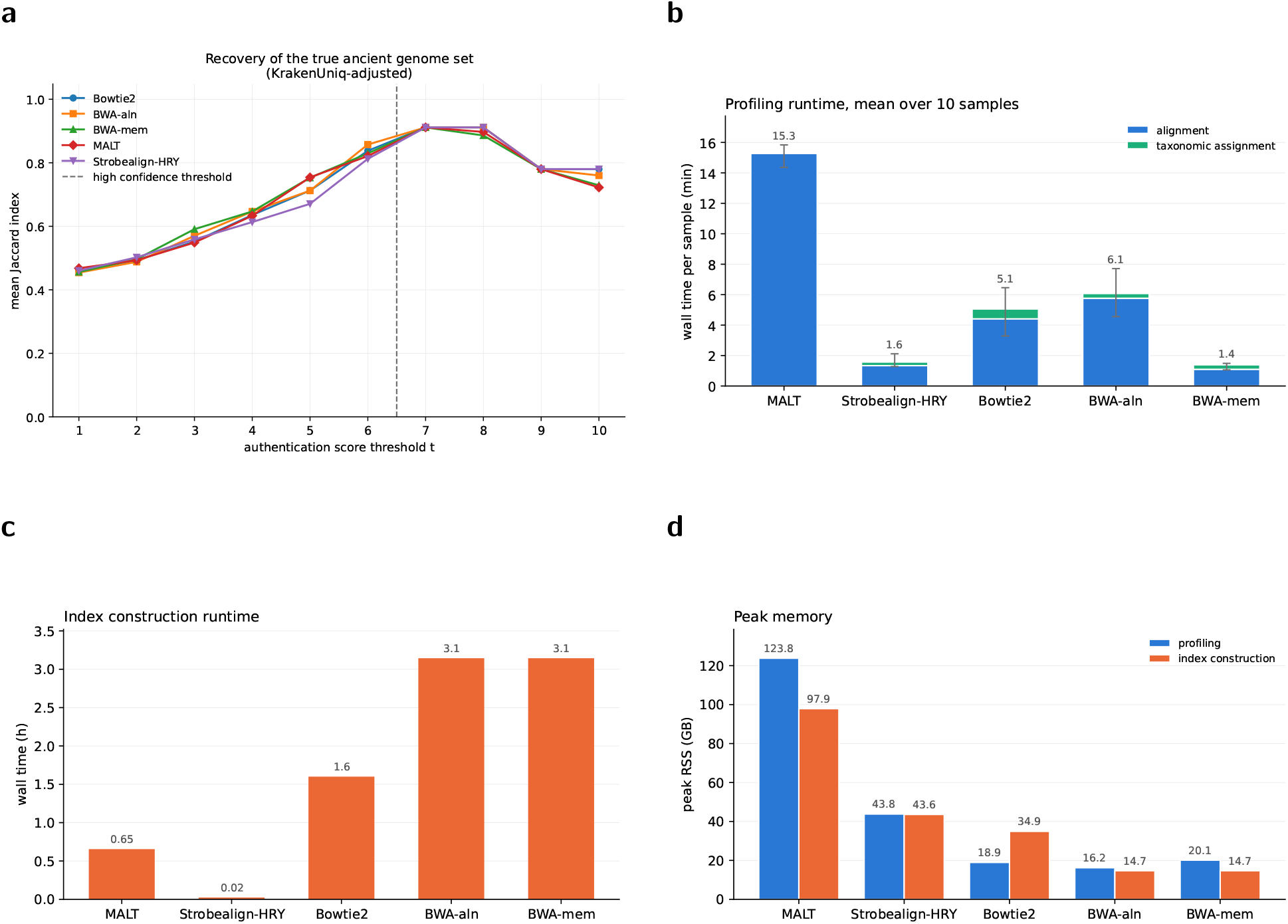
Effect of alignment method on post-screening ancient metagenomic profiling with aMeta. a. Mean Jaccard index between predicted and true ancient microbial sets across ten samples as the minimum authentication score varies. Taxa scoring at least the threshold are included in the predicted set. Both sets are restricted to taxa retained by KrakenUniq. The vertical dashed line represents the threshold after which no tool produces any false positives. b. Mean mapping and taxonomic-assignment runtime per sample, including ngsLCA for Strobealign-HRY, Bowtie2, BWA-aln, BWA-mem, and the built-in LCA assignment for MALT. c. One-time index construction runtime. d. Peak resident memory during mapping and index construction, shown separately.

At *t* = 7, all six false-negative calls belong to one species, Sorangium cellulosum. Its simulated genome has only 87.93% average nucleotide identity to the database reference used for authentication. All four methods give it a mean score deficit of five points relative to the maximum score of ten. This shared deficit is consistent with a limitation of reference representation rather than an aligner-specific failure (Supplementary Figure S3). The choice of aligner has a greater effect on computational cost than on recovery of the ancient community (Figure 3b). BWA-mem has the shortest mean runtime for alignment and taxonomic assignment, followed closely by Strobealign-HRY, and then by Bowtie2, BWA-aln, and MALT. These timings include alignment-rule processing and, where separate, the ngsLCA step. They exclude shared pre-screening, index construction, and downstream authentication. Strobealign-HRY also builds its index fastest, followed by MALT, Bowtie2, and both BWA versions (Figure 3c). For both mapping and index construction, Strobealign-HRY uses less peak memory than MALT, but more than Bowtie2 and both BWA versions (Figure 3d).

## Discussion

Hybrid RY-mer is a novel seed construct that combines multi-context seeds and R/Y encoding. When adapted into a genome aligner strobealign, it achieves noticeably higher alignment accuracy than other methods for ancient DNA alignment on the fruit fly and zymo dataset, with similar accuracy and highest precision on the CHM13 dataset. The improvement in alignment accuracy over other methods is particularly high in divergent samples. In terms of both mapping time and index construction, Strobealign-HRY is faster than BWA-aln, which is the most accurate tool for aDNA alignment [8]. While BWA-mem and Bowtie2 show better runtime than Strobealign-HRY, they are also less accurate for every dataset. Furthermore, by replacing the default aligner MALT within the aMETA pipeline with strobealign-HRY, mapping time is reduced by an order of magnitude, and peak memory is reduced by nearly a factor of 3.

A significant challenge in analysis of ancient samples is the limited amount of endogenous DNA that can be recovered from individual samples, as well as a finite amount of the ancient material [30, 31]. High mapping sensitivity of Strobealign-HRY can be utilized to obtain more information from limited ancient samples.

Deamination damage in aDNA molecules is both an obstacle to read mapping and a marker of ancient DNA authenticity. Uracil-DNA-glycosylase (UDG) is commonly used to remove uracil residues from DNA as part of sequencing library preparation for ancient samples [32, 33] to reduce deamination damage, however, the lack of deamination information makes the authentication of ancient samples more challenging. Partial UDG [34] treatment preserves the deamination signal while also reducing the fragment length. Since strobealign-HRY recovers more reads carrying deamination damage, it can be used as an alternative to UDG and partial UDG protocols in order to preserve scarce aDNA sequencing information.

### Future work

The main shortcoming of Strobealign-HRY is that it requires more memory than BWA and Bowtie2. The current memory layout is optimized for strobealign and strobemer seeds, and a more compact representation of hybrid RY-mers should reduce the memory cost. We also evaluated only a prefix layout for R/Y positions. Alternative patterns may result in a better balance between sensitivity and specificity, as shown for transition-constrained seeds [15]. Finally, the results of the metagenomic benchmark suggest that ancient species reconstruction is limited by the database selection and screening step, and is largely unaffected by alignment accuracy. Future work could investigate whether hybrid RY-mers can improve screening sensitivity for damaged sequences without substantially increasing false-positive candidates.

## Availability

Strobealign-HRY is implemented in Rust and publicly available at https://github.com/Itolstoganov/strobealign/releases/tag/hry. All data simulation and evaluation code used in this study is available at https://github.com/Itolstoganov/adna_bench.

## Acknowledgements

Kristoffer Sahlin was supported by the Swedish Research Council (SRC, Vetenskapsrådet) under Grant No. 2025-04304. Computational resources were provided by the National Academic Infrastructure for Supercom-puting in Sweden (NAISS), funded by the Swedish Research Council. We are also grateful to Zoé Pochon for her valuable feedback on the metagenomic assessment.

