## Supplementary Information for "Hybrid RY-mers increase aDNA alignment sensitivity"

### S1 Compared tools and evaluation criteria

Both strobealign configurations use commit `4be2dcc`. Strobealign-HRY uses `-adna -ry-len 8 -M 500 -ssw`, and Strobealign-DK uses `-l 1 -u 1 -M 500`. BWA-aln uses `-n 0.01 -o 2 -l 1024`, BWA-MEM uses `-r 2.5`, and Bowtie2 uses `-local -very-sensitive-local`. All mapping runs use four threads. We vary k from 13 to 21 in steps of two for Strobealign and BWA-MEM; both strobealign configurations use s = k - 2. Figure 2 uses k = 17 and s = 15 for strobealign and a minimum seed length of 17 for BWA-MEM. The hybrid RY-mer length parameter is fixed at 8 throughout the Strobealign-HRY runs.

Strobealign timings cover the reported mapping phase. Separate index construction is excluded, as is BWA-aln's subsequent SAM generation step. Peak memory is the maximum resident set size of the timed aligner process.

### S2 Individual aMeta authentication criteria results

The individual authentication criteria show a similar pattern of missed signals across methods, although their false-positive counts differ (Supplementary Figure S4). All methods fail the evenness-of-coverage criterion for 16 ancient taxon-sample combinations, and the edit-distance criterion for all reads for six. The table evaluates each criterion separately on ancient taxa present in a sample and candidate taxa absent from that sample, after restricting both groups to KrakenUniq candidates. Modern taxa present in the sample are excluded. A taxon-sample combination can fail multiple criteria, so these counts do not represent false-negative ancient calls based on the combined authentication score.

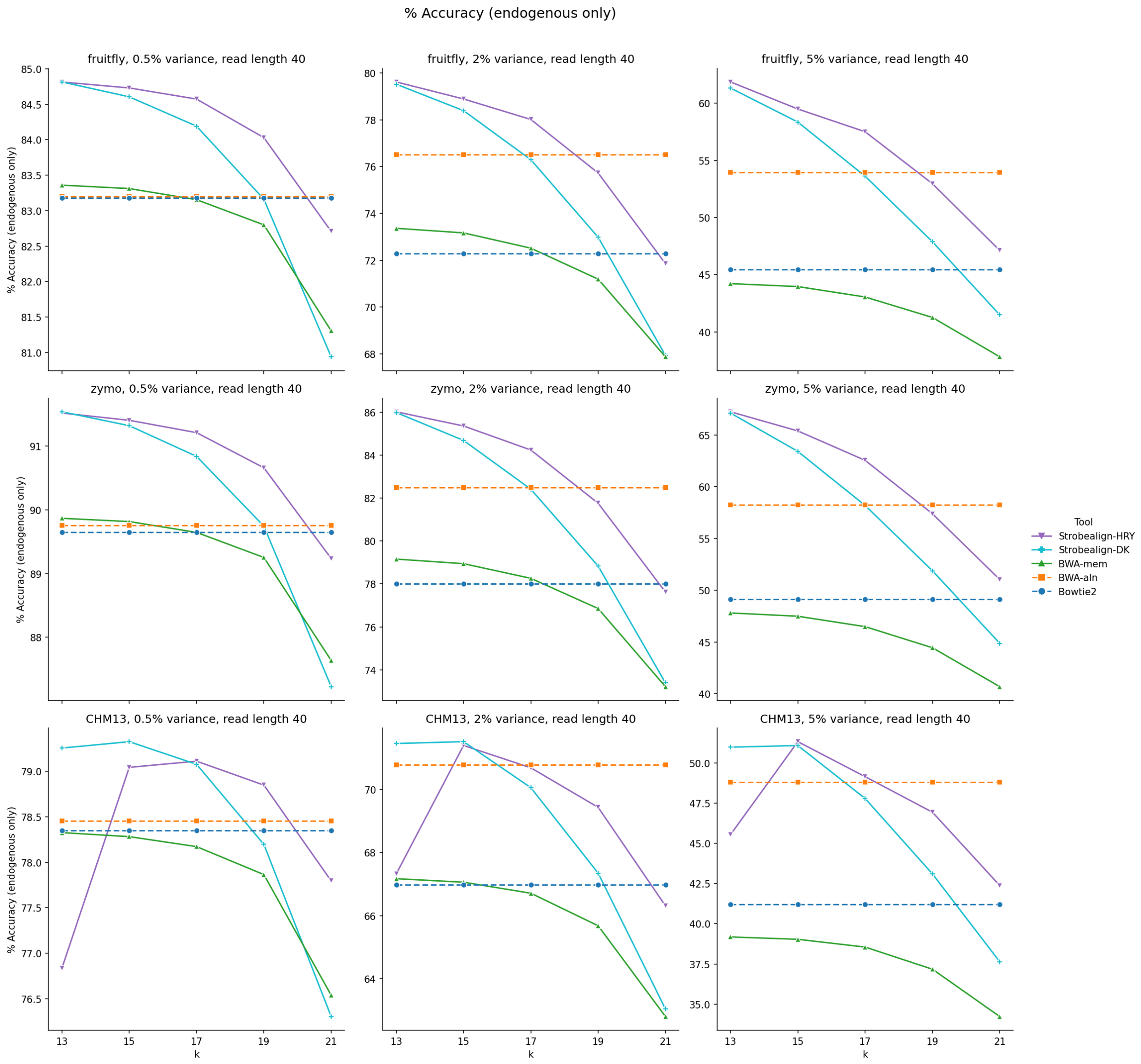


**Supplementary Figure S1:** Percentage of correctly mapped reads for fruit fly (top), Zymo (middle), and CHM13 (bottom) datasets. Values are shown across different values of parameter $k$ for both Strobealign versions and BWA-mem.

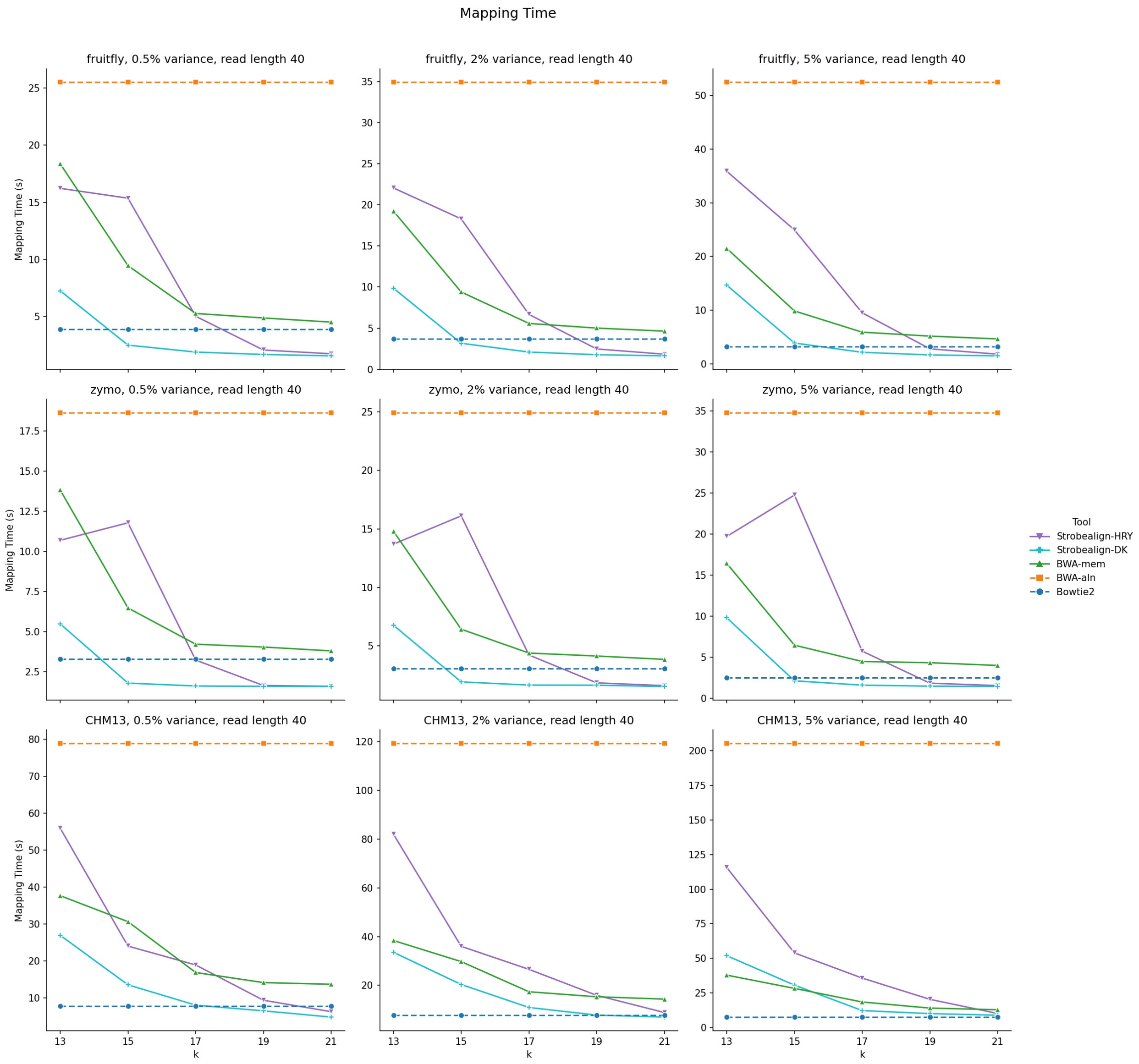


**Supplementary Figure S2:** Total mapping time (in seconds) for fruit fly (top), Zymo (middle), and CHM13 (bottom) datasets. Values are shown across different values of parameter $k$ for both Strobealign versions and BWA-mem.

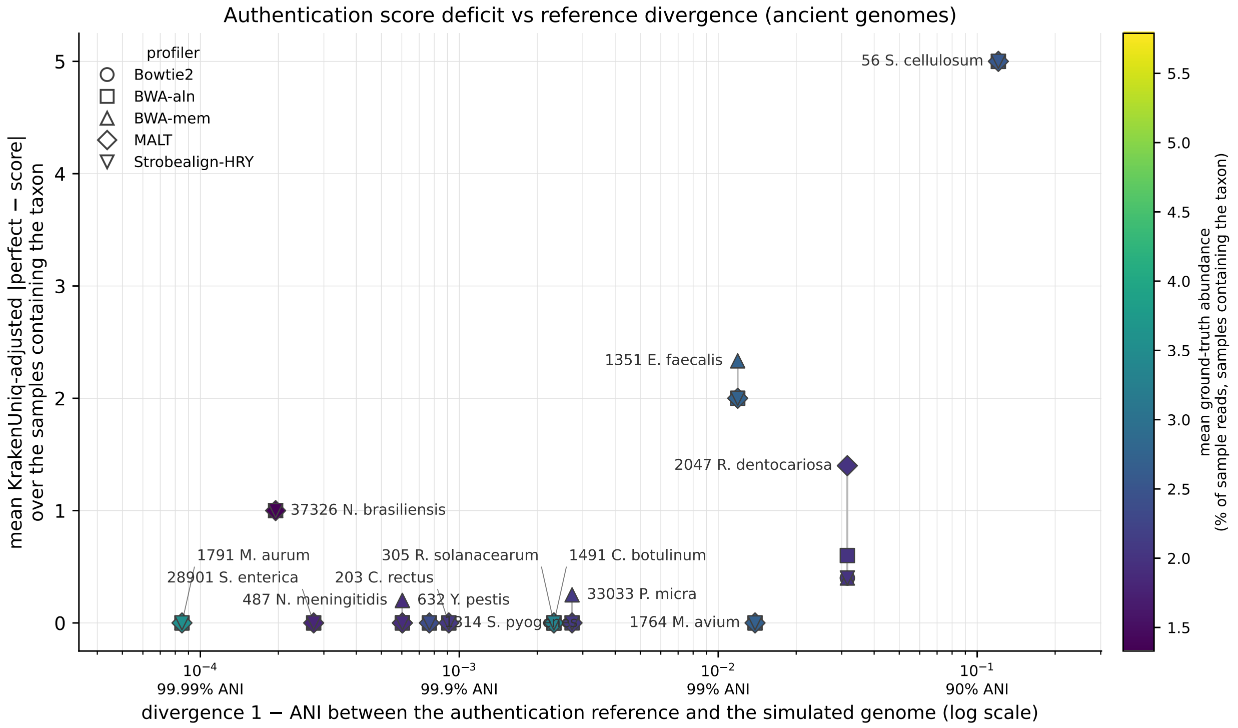


**Supplementary Figure S3:** KrakenUniq-adjusted mean authentication-score deficit versus divergence between each simulated ancient genome and its authentication reference, measured as 1 - ANI on a logarithmic scale.

Authentication criteria results (ancient genomes)

| | | Bowtie2 | | BWA-aln | | BWA-mem | | MALT | | Strobealign-HRY | |
|---|---|---|---|---|---|---|---|---|---|---|---|
| **Criterion** | **Points** | **FP** | **FN** | **FP** | **FN** | **FP** | **FN** | **FP** | **FN** | **FP** | **FN** |
| damage C>T_1 | 1 | 8 | 0 | 8 | 0 | 5 | 0 | 13 | 0 | 9 | 0 |
| damage G>A_20 | 1 | 11 | 0 | 6 | 0 | 5 | 0 | 14 | 0 | 12 | 0 |
| evenness of coverage | 2 | 7 | 16 | 4 | 16 | 7 | 16 | 15 | 16 | 10 | 16 |
| edit distance (all) | 1 | 6 | 6 | 6 | 6 | 6 | 7 | 6 | 6 | 6 | 6 |
| edit distance (ancient) | 1 | 0 | 6 | 0 | 7 | 1 | 10 | 2 | 6 | 1 | 6 |
| PMD | 1 | 35 | 0 | 28 | 0 | 31 | 0 | 44 | 0 | 34 | 0 |
| read length | 1 | 10 | 0 | 28 | 0 | 10 | 0 | 5 | 0 | 7 | 0 |
| depth of coverage | 1 | 34 | 0 | 33 | 0 | 35 | 0 | 39 | 0 | 34 | 0 |
| ANI | 1 | 8 | 6 | 9 | 6 | 9 | 6 | 7 | 11 | 7 | 6 |
| **total** | **10** | **119** | **34** | **122** | **35** | **109** | **39** | **145** | **39** | **120** | **34** |

**Supplementary Figure S4:** KrakenUniq-adjusted false-positive and false-negative counts for individual aMeta authentication criteria. The evaluation includes ancient taxa present in each sample and candidate taxa absent from that sample; modern taxa present in the sample are excluded. A false positive means that a criterion is satisfied for an absent taxon, and a false negative means that an ancient taxon fails a criterion. Counts are pooled across ten samples. The points column gives each criterion's contribution to the ten-point authentication score. A taxon-sample combination may contribute to multiple criterion-level errors.
